# Inactivation analysis of SARS-CoV-2 by specimen transport media, nucleic acid extraction reagents, detergents and fixatives

**DOI:** 10.1101/2020.07.08.194613

**Authors:** Stephen R. Welch, Katherine A. Davies, Hubert Buczkowski, Nipunadi Hettiarachchi, Nicole Green, Ulrike Arnold, Matthew Jones, Matthew J. Hannah, Reah Evans, Christopher Burton, Jane E. Burton, Malcolm Guiver, Patricia A. Cane, Neil Woodford, Christine B. Bruce, Allen D. G. Roberts, Marian J. Killip

## Abstract

The COVID-19 pandemic has necessitated a rapid multi-faceted response by the scientific community, bringing researchers, health officials and industry together to address the ongoing public health emergency. To meet this challenge, participants need an informed approach for working safely with the etiological agent, the novel human coronavirus SARS-CoV-2. Work with infectious SARS-CoV-2 is currently restricted to high-containment laboratories, but material can be handled at a lower containment level after inactivation. Given the wide array of inactivation reagents that are being used in laboratories during this pandemic, it is vital that their effectiveness is thoroughly investigated. Here, we evaluated a total of 23 commercial reagents designed for clinical sample transportation, nucleic acid extraction and virus inactivation for their ability to inactivate SARS-CoV-2, as well as seven other common chemicals including detergents and fixatives. As part of this study, we have also tested five filtration matrices for their effectiveness at removing the cytotoxic elements of each reagent, permitting accurate determination of levels of infectious virus remaining following treatment. In addition to providing critical data informing inactivation methods and risk assessments for diagnostic and research laboratories working with SARS-CoV-2, these data provide a framework for other laboratories to validate their inactivation processes and to guide similar studies for other pathogens.

## 1. Introduction

Infection with the novel human betacoronavirus SARS-CoV-2 can cause a severe or fatal respiratory disease, termed COVID-19 (1–3). As the COVID-19 pandemic has developed, millions of clinical samples have been collected for diagnostic evaluation. SARS-CoV-2 has been classified as a Hazard Group 3 pathogen in the UK, and as such, deliberate work with the virus must be carried out in high containment laboratories (containment level 3 (CL3) in the UK) with associated facility, equipment and staffing restrictions. Guidance from Public Health England (PHE) and the World Health Organization (WHO) enable non-propagative testing of clinical specimens to be carried out at the lower CL2, with the requirement that all non-inactivated material is handled within a microbiological safety cabinet (MSC) and that the process has been suitably and sufficiently risk assessed (4, 5). Guidance from the U.S. Centers for Disease Control and Prevention requires that specimens must be inactivated (e.g. in nucleic acid extraction buffer) before handling at biosafety level 2 (BSL-2) (6). To allow safe movement of clinical samples from CL3/BSL-3 laboratories to CL2/BSL-2, virus inactivation procedures should be validated, and formal validation of inactivation protocols are often an operational requirement for clinical and research laboratories handling SARS-CoV-2.

Efficacy of virus inactivation depends on numerous factors, including the nature and concentration of pathogen, sample matrix, concentration of inactivation agent/s and contact time. To date, there are limited data on efficacy of SARS-CoV-2-specific inactivation approaches in the scientific literature and risk assessments have largely been based upon inactivation information for genetically related coronaviruses. Previous studies have found that treatment with heat, chemical inactivants, ultraviolet light, gamma irradiation and a variety of detergents are effective at inactivating SARS-CoV-1 and Middle East Respiratory Syndrome coronavirus (MERS-CoV), other high-consequence human coronaviruses (7–13). However, limited validation data exist for coronavirus inactivation by commercial sample transport media and molecular extraction lysis buffers used in steps prior to nucleic acid extraction for diagnostic testing. Furthermore, the precise composition of many commercial reagents is proprietary, preventing ingredient-based inference of inactivation efficacy between reagents. Some limited preliminary data on SARS-CoV-2 inactivation are available (14–19), but given the current level of diagnostic and research activities, there is an urgent need to comprehensively investigate SARS-CoV-2-specific inactivation efficacy of available methods to support safe virus handling.

An important consideration in inactivation assays is cytotoxicity, a typical effect of many chemical inactivants. To mitigate cytotoxic effects, the inactivation agent needs to be either diluted out or removed from treated samples prior to testing for infectious virus. Each of these methods for addressing cytotoxicity present their own challenges. Sample dilution requires the use of high titer stocks of virus (e.g. >10^8^ PFU/mL) to be able to demonstrate a significant titer reduction and reduces recovery of low level residual virus from treated samples, making it difficult or impossible to distinguish complete from incomplete virus inactivation. In contrast, methods for purification of virus away from cytotoxic components in treated samples may also remove virus or affect virus viability. Accurate quantification of remaining infectious virus ideally requires complete removal of cytotoxicity without compromising assay sensitivity, which needs careful consideration of reagent and purification processes prior to performing inactivation tests.

Here, we describe optimal methods for the removal of cytotoxicity from samples treated with commercial reagents, detergents and fixatives. These data were then used in evaluations of the effectiveness of these chemicals for inactivating SARS-CoV-2. This work, applicable to both diagnostic and research laboratories, provides invaluable information for public health and basic research responses to the COVID-19 pandemic by supporting safe approaches for collection, transport, extraction and analysis of SARS-CoV-2 samples. Furthermore, our studies investigating purification of a wide range of cytotoxic chemicals are highly applicable to inactivation studies for other viruses, thereby supporting rapid generation of inactivation data for known and novel viral pathogens.

## 2. Materials and Methods

### 2.1. Cells and virus

Vero E6 cells (Vero C1008; ATCC CRL-1586) were cultured in modified Eagle’s minimum essential medium (MEM) supplemented with 10% (v/v) fetal calf serum (FCS). Virus used was SARS-CoV-2 strain hCOV-19/England/2/2020, isolated by PHE from the first patient cluster in the UK on 29/01/2020. This virus was obtained at passage 1 and used for inactivation studies at passage 2 or 3. All infectious work carried out using an MSCIII in a CL3 laboratory. Working virus stocks were generated by infecting Vero E6 cells at a multiplicity of infection (MOI) of 0.001, in the presence of 5% FCS. Cell culture supernatants were collected 72 hours post infection, clarified for 10 mins at 3000 × g, aliquoted and stored at −80°C until required. Viral titers were calculated by either plaque assay or 50% tissue culture infectious dose (TCID50). For plaque assays, 24-well plates were seeded the day before the assay (1.5 × 10^5^ cells/well in MEM/10%FCS). Ten-fold dilutions of virus stock were inoculated onto plates (100μL per well), inoculated at room temperature for 1 hour then overlaid with 1.5% medium viscosity carboxymethylcellulose (Sigma-Aldrich) and incubated at 37°C/5% CO_2_ for 3 days. For TCID50s, ten-fold dilutions of virus stock (25μL) were plated onto 96-well plates containing Vero E6 cell suspension (2.5 × 10^4^ cells/well in 100μl MEM/5%FCS) and incubated at 37°C/5% CO_2_ for 5-7 days. Plates were fixed with 4% (v/v) formaldehyde/PBS, and stained with 0.2% (v/v) crystal violet/water TCID50 titers were determined by the Spearman-Kärber method (20, 21).

### 2.2. Reagents and chemicals used for SARS-CoV-2 inactivation

The commercial reagents evaluated in this study, along with their compositions (if known) and manufacturers’ instructions for use (if provided) are given in Supplementary Table 1. Specimen transport reagents tested were: Sigma Molecular Transport Medium (MM, Medical Wire); eNAT (Copan); Primestore Molecular Transport Medium (MTM, Longhorn Vaccines and Diagnostics); Cobas PCR Media (Roche); Aptima Specimen Transport Medium (Hologic); DNA/RNA Shield, (Zymo Research); guanidine hydrochloride (GCHl) and guanidine thiocyanate (GITC) buffers containing Triton X-100 (both Oxoid/Thermo Fisher); Virus Transport and Preservation Medium Inactivated (BioComma). Molecular extraction reagents tested were: AVL, RLT, ATL, and AL (all Qiagen); MagNA Pure external lysis buffer, and Cobas Omni LYS used for on-board lysis by Cobas extraction platforms (Roche); Viral PCR Sample Solution (VPSS) and Lysis Buffer (both E&O Laboratories); NeuMoDx Lysis Buffer (NeuMoDx Molecular); Samba II SCoV lysis buffer (Diagnostics for the Real World); NucliSENS lysis buffer (Biomerieux); Panther Fusion Specimen Lysis Tubes (Hologic); and an in-house extraction buffer containing guanidine thiocyanate and Triton X-100 (PHE Media Services). Detergents tested were: Tween 20, Triton X-100 and NP-40 Surfact-Amps Detergent Solutions (all Thermo Scientific), and UltraPure SDS 10% solution (Invitrogen). Other reagents assessed include: polyhexamethylene biguanide (PHMB, Blueberry Therapeutics); Formaldehyde and Glutaraldehyde (TAAB); and Ethanol and Methanol (Fisher Scientific).

### 2.3. Removal of reagent cytotoxicity

Specimen transport tube reagents were assessed undiluted unless otherwise indicated. For testing of molecular extraction reagents, mock samples were generated by diluting reagent in PBS at ratios given in manufacturer’s instructions. Detergents, fixatives and solvents were all assessed at the indicated concentrations. All methods were evaluated in a spin column format, for ease of sample processing within the high containment laboratory. Pierce Detergent Removal Spin Columns (0.5mL, Thermo Scientific), Microspin Sephacryl S400HR (GE Healthcare), and Amicon Ultra-0.5mL 50KDa centrifugal filters (Merck Millipore) were prepared according to manufacturer’s instructions. Sephadex LH-20 (GE Healthcare) and Bio-Beads SM2 resin (Bio-Rad) were suspended in PBS and poured into empty 0.8mL Pierce centrifuge columns (Thermo Scientific), and centrifuged for one min at 1000 × g to remove PBS immediately before use. For all matrices aside from the Amicon Ultra columns, 100μl of treated sample was added to each spin column, incubated for two mins at room temperature, then eluted by centrifugation at 1,000 × g for two mins. For Amicon Ultra filters, 500μl of sample was added, centrifuged at 14,000 × g for 10 mins, followed by three washes with 500μl PBS. Sample was then collected by resuspending contents of the filtration device with 500μl PBS. To assess remaining cytotoxicity, a two-fold dilution series of treated filtered sample was prepared in PBS, and 6.5μl of each dilution transferred in triplicate to 384-well plates containing Vero E6 cells (6.25 × 10^3^ cells/well in 25μl MEM/5%FCS) and incubated overnight. Cell viability was determined by CellTiter Aqueous One Solution Cell Proliferation Assay (Promega) according to manufacturer’s instructions. Normalized values of absorbance (relative to untreated cells) were used to fit a 4-parameter equation to semilog plots of the concentration-response data, and to interpolate the concentration that resulted in 80% cell viability (CC20) in reagent treated cells. All analyses were performed using GraphPad Prism 8 (v8.4.1, GraphPad Software).

### 2.4. SARS-CoV-2 inactivation

For commercial products, virus preparations (tissue culture fluid, titers ranging from 1 × 10^6^ to 1 × 10^8^ PFU/ml) were treated in triplicate with reagents at concentrations and for contact times recommended in the manufacturers’ instructions for use, where available, or for concentrations and times specifically requested by testing laboratories. Where a range of concentrations was given by the manufacturer, the lowest ratio of product to sample was tested (i.e. lowest recommended concentration of test product). Specimen transport tube reagents were tested using a ratio of one volume of tissue culture fluid to ten volumes of reagent, unless a volume ratio of sample fluid to reagent was specified by the manufacturer. Detergents, fixatives and solvents were tested at the indicated concentrations for the indicated times. For testing of alternative sample types, virus was spiked into the indicated sample matrix at a ratio of 1:9, then treated with test reagents as above. All experiments included triplicate control mock-treated samples with an equivalent volume of PBS in place of test reagent. Immediately following the required contact time, 1mL of treated sample was processed using the appropriately selected filtration matrix. Reagent removal for inactivation testing was carried out in a larger spin column format using Pierce 4mL Detergent Removal Spin Columns (Thermo Fisher), or by filling empty Pierce 10mL capacity centrifuge columns (Thermo Fisher) with SM2 Bio-Beads, Sephacryl S-400HR or Sephadex LH-20 to give 4mL packed beads/resin. For purification using Amicon filters, 2 × 500μl samples were purified using two centrifugal filters by the method previously described, then pooled together. For formaldehyde and formaldehyde with glutaraldehyde removal, one filter was used with 1 × 500μl sample volume, resuspended after processing in 500μl PBS, and added to 400ul MEM/5% FBS. For inactivation of infected monolayers, 12.5 cm^2^ flasks of Vero E6 cells (2.5 × 10^6^ cells/flask in 2.5mL MEM/5% FBS) were infected at MOI 0.001 and incubated at 37°C/5% CO_2_ for 24 hours. Supernatant was removed, and cells fixed using 5mL of formaldehyde, or formaldehyde and glutaraldehyde at room temperature for 15 or 60 mins. The fixative was removed, and monolayers washed three times with PBS before scraping cells into 1mL MEM/5% FBS and sonicated (3 × 10 second on,10 seconds off at 100% power and amplitude) using a UP200St with VialTweeter attachment (Hielscher Ultrasound Technology). Supernatants were clarified by centrifuging at 3000 × g for 10 mins.

### 2.5. SARS-CoV-2 quantification and titer reduction evaluation

Virus present in treated and purified, or mock-treated and purified, samples was quantified by either TCID50 or plaque assay. As additional assay controls, unfiltered mock-treated sample was titrated to determine virus loss during filtration, and filtered test-reagent only (no virus) sample titrated to determine residual test buffer cytotoxicity. For TCID50 assays, neat to 10^−7^ ten-fold dilutions were prepared, and for plaque assays, neat to 10^−5^ ten-fold dilutions were prepared, both in MEM/5% FCS. TCID50 titers were determined by the Spearman-Kärber method (20, 21). Conditions where low levels of virus were detected such that TCID50 could not be calculated by Spearman-Kärber, TCID50 was calculated the Taylor method (22). Where no virus was detectable, values are given as less than or equal to the Taylor-derived TCID50 titer given by a single virus positive well at the lowest dilution where no cytotoxicity was observed. Titer reduction was calculated by subtracting the mean logarithmic virus titer for test-buffer-treated, purified conditions from the mean logarithmic virus titer for the PBS-treated, purified condition, with standard errors calculated according to (22).

### 2.6. Serial passages of treated samples

In parallel to virus quantification, 12.5 cm^2^ flasks of Vero E6 cells (6.25 × 10^4^ cells/flask in 2.5mL MEM/5% FBS) were inoculated with either 500μl or 50μl of treated filtered sample. Flasks were examined for cytopathic effect (CPE) and 500μl culture medium from each flask was used to inoculate new 12.5 cm^2^ flasks of Vero E6 cells after seven days. If no CPE was observed, this process was continued for up to four serial passages. For the duration of the passage series, a flask of untreated cells was included as a control for cross-contamination between flasks, and a SARS-CoV-2 infected control was included to ensure suitable conditions for virus propagation. To distinguish CPE from any residual cytotoxicity associated with test reagents, samples of cell culture medium were taken from each flask at the beginning and end of each passage. Nucleic acid was extracted from cell culture media manually using a QIAamp Viral RNA Mini Kit (QIAGEN) or using NucliSENS easyMAG or EMAG platforms (both BioMérieux). Viral RNA levels were quantified by quantitative reverse-transcriptase PCR (qRT-PCR) specific for the SARS-CoV-2 E gene (23) using TaqMan Fast 1-Step Master Mix (Applied Biosystems) on a 7500 Fast Real-Time PCR System (Applied Biosystems). A positive result for virus amplification was recorded if effects on the monolayer consistent with CPE and a decrease in Ct across the course of a passage were observed.

## 3. Results

### 3.1. Reagent filtration optimization to minimize cytotoxicity and maximum virus recovery

Prior to evaluating their effectiveness at inactivating SARS-CoV-2, we investigated the cytotoxicity of each reagent before and after filtration though one of five matrices: Sephadex LH-20, Sephacryl S400HR, Amicon Ultra 50kDa molecular weight cut-off centrifugal filters, Pierce detergent removal spin columns (DRSC), and Bio-Beads SM2 nonpolar polystyrene adsorbents. Reagents were diluted with PBS to the working concentrations recommended by the manufacturer (for commercial sample transport and molecular extraction reagents), or the indicated concentrations (for all other chemicals), followed by a single reagent removal step with each filtration matrix. Dilution series of filtered and unfiltered samples were generated to determine concentration-dependent cytotoxicity, from which the CC20 value for each combination of reagent and filtration method were interpolated (Supplementary Figure 1). CC20 was chosen as, at this concentration, cells retain 80% viability and enable distinction of active SARS-CoV-2 replication by visualisation of CPE in the monolayer. Table 1 shows the dilution factor of reagent-treated sample required to achieve the CC20 after filtration, with <1 indicating complete removal of cytotoxicity. These data were used to determine the relative cytotoxicity removed by one filtration step for each combination of reagent and matrix (Figure 1A).

**Figure 1:**
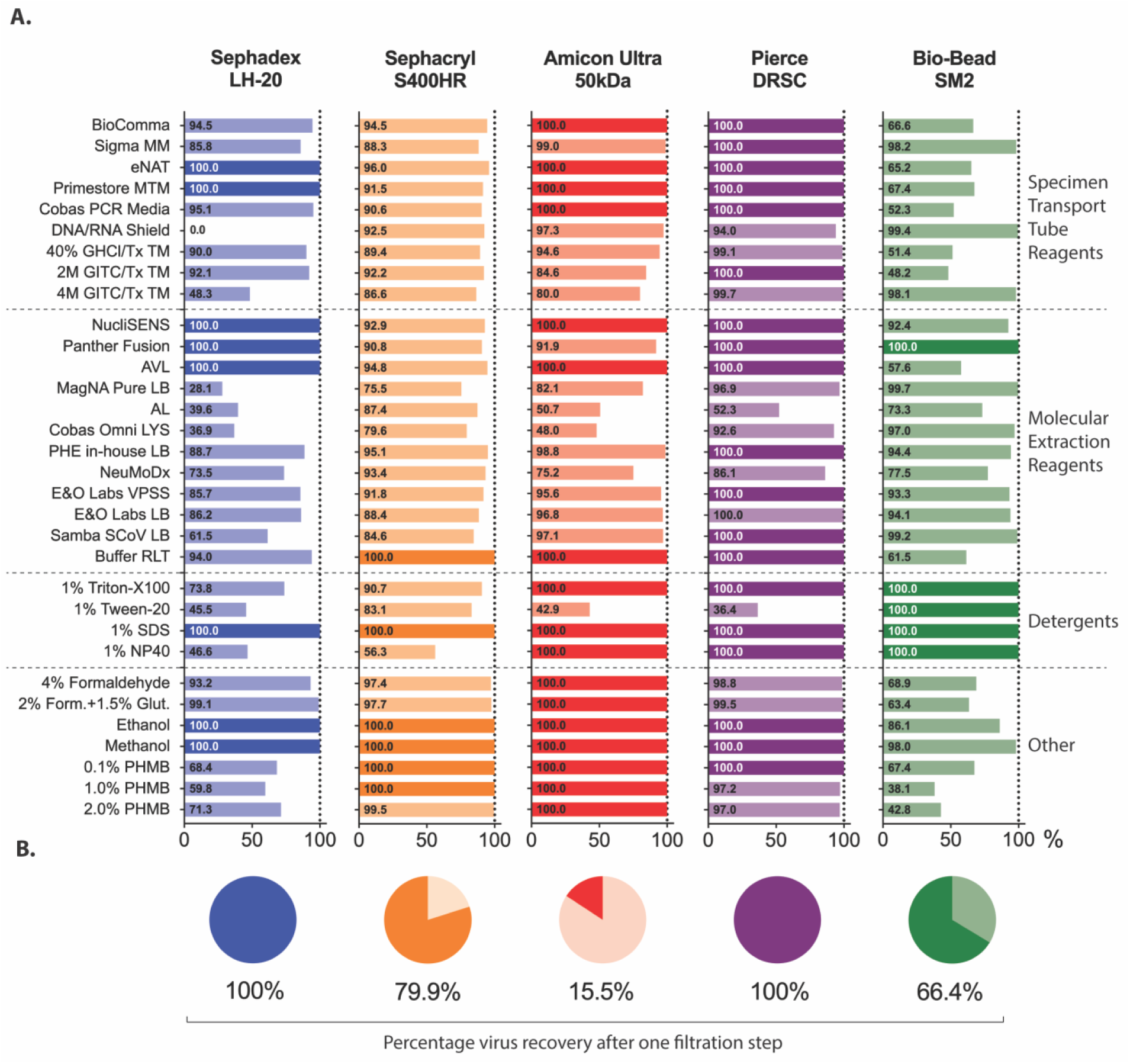
Effectiveness of five filtration matrices at removing cytotoxicity. **(A)** SARS-CoV-2 virus in clarified cell culture supernatant was treated with indicated reagent for 2mins at room temperature before being purified through one of 5 filtration matrices: Sephadex LH-20 (blue); Sephacryl S400HR (orange); Amicon Ultra 50kDa molecular weight cut off (red); Pierce detergent removal spin columns (DRSC) (purple); or Bio-Bead SM2 (green). Values indicate the percentage toxicity removal after one purification cycle relative to unpurified samples (based on CC20 values – for more details see Table 1). **(B)** Percentage of input virus remaining in eluate after one purification cycle through each filtration matrix. GHCl - guanidine hydrochloride; GITC - guanidinium isothiocyanate; Tx – Triton X-100; PHMB - polyhexamethylene biguanide; SDS - sodium dodecyl sulfate; NP40 - nonyl phenoxypolyethoxylethanol. LB – lysis buffer; TM – transport medium

**Table 1:**
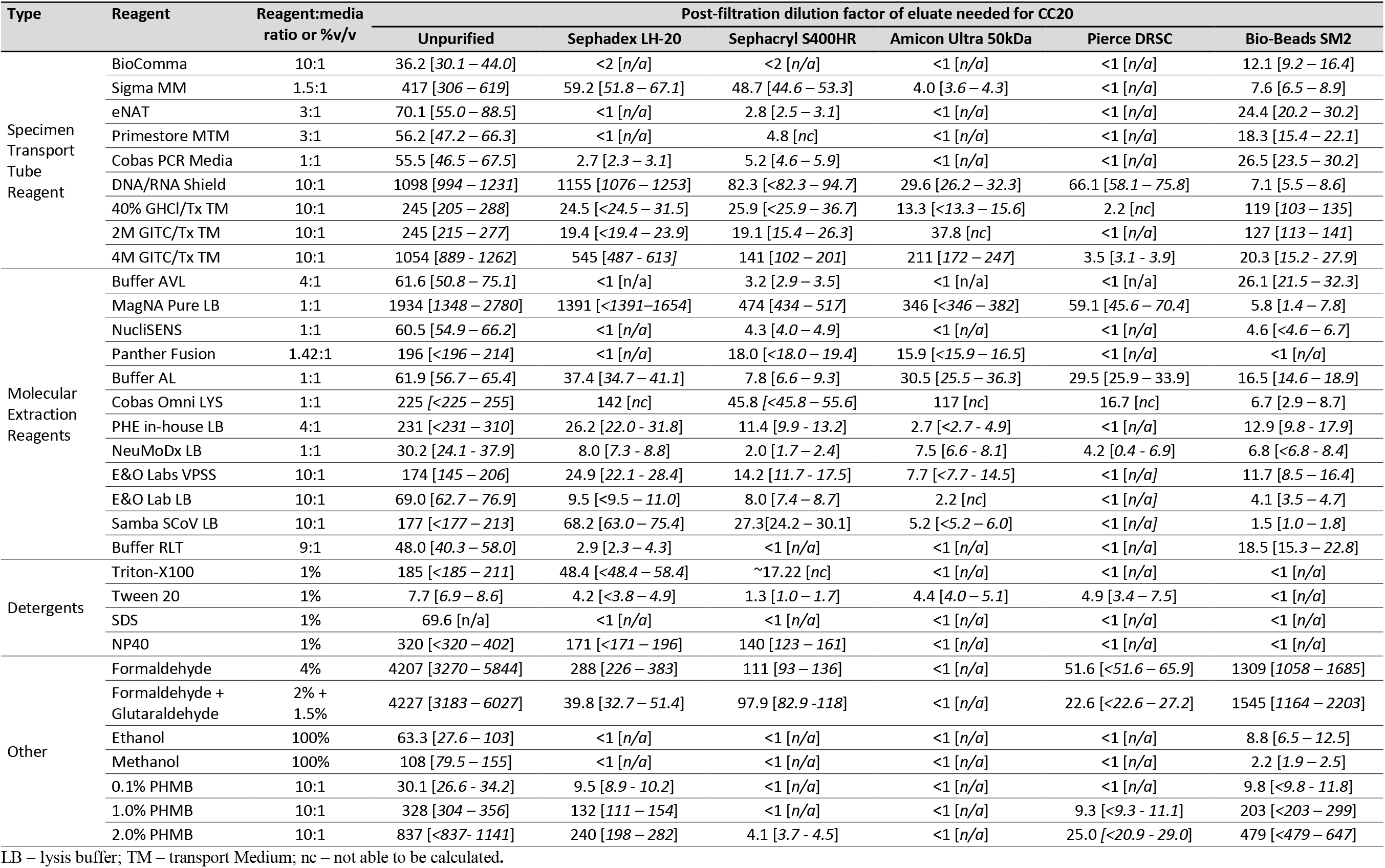
Purification of reagents: Values [95% CI] represent the dilution factor required after one purification process to achieve the CC20 concentration [95% CI].

All unfiltered reagents tested here were cytotoxic, but the degree of cytotoxicity varied considerably as did the optimal filtration matrix for each reagent. The detergent Tween 20 used at 1% concentration was the least cytotoxic unfiltered, only requiring a dilution factor of 7.7 to reach the CC20, although only the Bio-Bead SM2 filters were effective at removing all cytotoxicity. The chemical fixative combination of 2% formaldehyde plus 1.5% glutaraldehyde was the most cytotoxic unfiltered, requiring a dilution of over 4000 to reach the CC20, with only the Amicon Ultra columns able to remove 100% of the cytotoxicity. However, for the majority reagents (27/34) tested, filtration through at least one matrix type removed 100% of cytotoxicity allowing neat eluate to be used directly in cell culture without further dilution. There were several exceptions to this: DNA/RNA shield (maximum 99.4% cytotoxicity removal using SM2); 40% GHCl (99.1% using Pierce DRSC); 4M GITC (99.7% using Pierce DRSC); MagNA Pure (99.7% using SM2); AL buffer (87.4% using S400HR); Cobas Omni LYS (97.0% using SM2); and NeuMoDx (93.4% using S400HR). For these reagents, filtered eluate was still cytotoxic when used undiluted in cell culture. However, CC20 values indicated that this remaining cytotoxicity would be removed by first or second (10^−1^ – 10^−2^) dilutions in the TCID50 assay allowing evaluation of titer reduction using these reagents with the caveat that the effective assay limit of detection (LOD) would be higher. Passing treated samples through more than one column, or increasing the depth of the resin/bead bed within the spin column can also improve cytotoxicity removal for some reagents (unpublished data).

In addition to cytotoxicity removal, a successful filtration method must also purify virus without adversely affecting titer or integrity. We therefore assessed SARS-CoV-2 recovery after each filtration method. Using an input titer of 1.35 × 10^6^ TCID50/mL, triplicate purifications of virus through Sephadex LH-20 or Pierce detergent removal spin columns resulted in recovery of 100% of input virus (Figure 1B). In contrast, the recoverable titer after one filtration through Amicon Ultra filters was 2.13 × 10^5^ TCID50/mL, an 85% reduction from input. Purification with S400HR and Bio-Beads SM2 matrices resulted in recoverable titers of 1.08 × 10^6^ TCID50/mL and 8.99 × 10^5^ TCID50/mL, a loss of 30% and 35% of input virus, respectively.

### 3.2. SARS-CoV-2 inactivation by specimen transport and molecular extraction reagents

Specimen transport tubes are designed to inactivate microorganisms present in clinical specimens prior to sample transport, while preserving the integrity of nucleic acids for molecular testing. If effective, these products have the potential to streamline SARS-CoV-2 diagnostic processing in testing laboratories by eliminating the requirement for CL3 processing or, for activities derogated to CL2, permitting processing outside an MSC. The BS EN 14476 standard requires demonstration of a >4 log_10_ titer reduction for virucidal suspension tests (22), and we were able to demonstrate a ≥4 log10 TCID50 titer reduction for all specimen transport media evaluated in a tissue culture fluid matrix (Table 2). However, infectious virus remained recoverable in treated samples after inactivation with most reagents tested (by either TCID50 or blind passage). The exceptions to this were PrimeStore MTM and 4M GITC, from which no residual virus was detectable by either TCID50 or by the passaging of treated purified sample. While several contact times were evaluated for all these reagents, length of contact time had no effect on either the level of virus titer reduction or whether virus remained detectable upon passage.

**Table 2:**
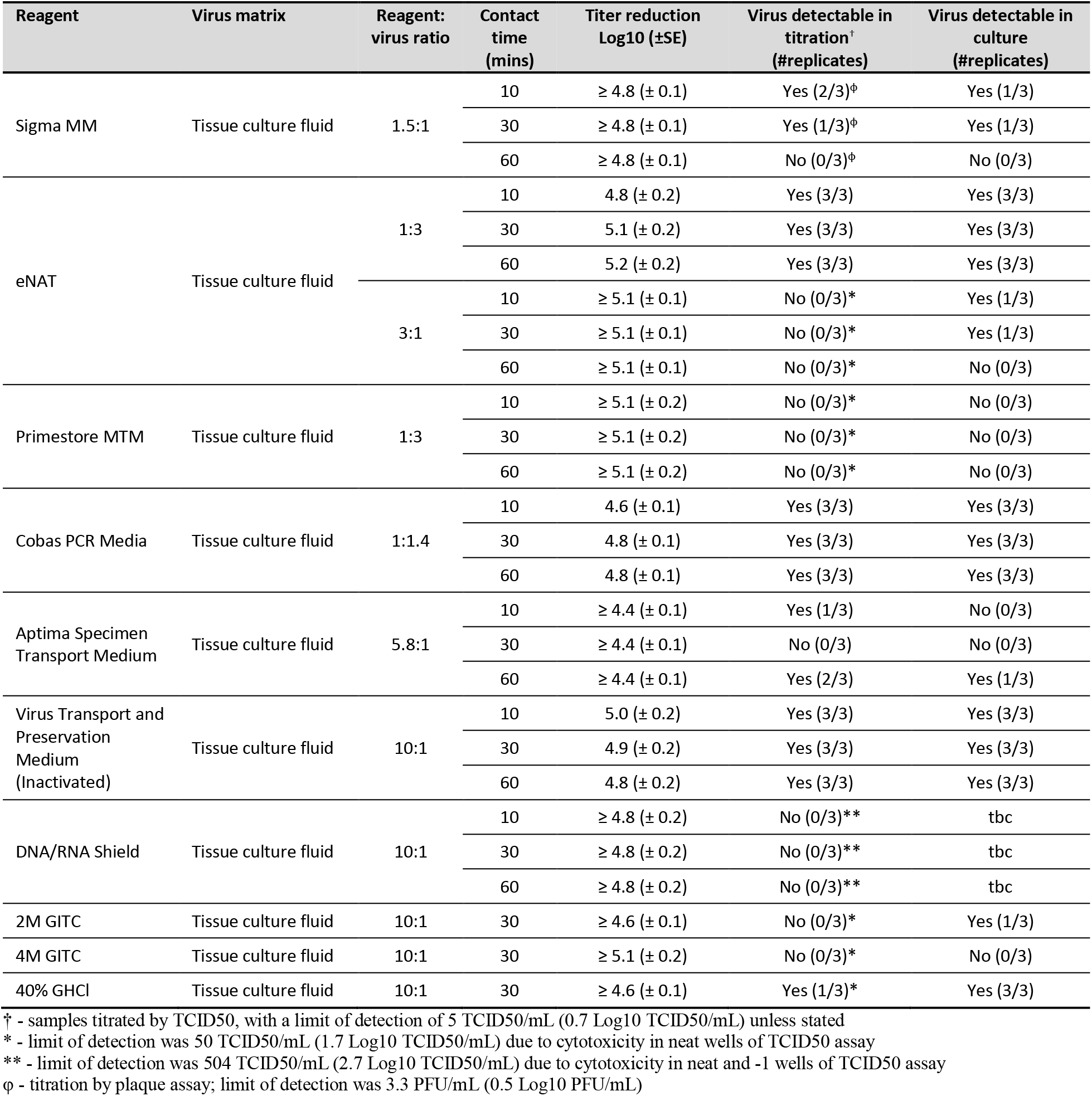
Virus inactivation by specimen transport tube reagents

We also sought to inform sample processing by examining inactivation by molecular extraction lysis buffers used in several manual and automated extraction protocols within SARS-CoV-2 diagnostic and research laboratories. We could demonstrate a ≥4 log10 reduction in TCID50 titer for all but two molecular extraction reagents when evaluated using tissue culture fluid (Table 3). The exceptions to this were AL and Cobas Omni LYS, where remaining cytotoxicity in the filtered eluate increased the TCID50 LOD to a level such that the maximum calculable titer reductions were ≥3.5 and ≥3.9 log10 TCID50s, respectively. However, given no virus was detected at any passage it is likely that infectious virus was effectively inactivated by these two reagents. For reagents tested with multiple contact times (NucliSENS, Panther Fusion), shorter times (10 mins) were as effective at reducing virus titers as longer contact times. Most reagents reduced viral titers to around the TCID50 assay LOD, indicating that any remaining virus post treatment was present only at very low titers (<10 TCID50/mL), but higher levels of virus were recoverable from samples treated with some extraction buffers. For NeuMoDx lysis buffer, although titers were reduced by ≥4 log10 TCID50s, an average of 91 (±38) TCID50/mL remained detectable. Similarly, Buffer AVL reduced virus titers by 5.1 log10 TCID50s, but after treatment virus was detectable in all treated samples replicates (average 54 (±18) TCID50/mL). However, addition of four sample volumes of absolute ethanol following a 10 minute contact time with AVL (the next step in the QIAGEN Viral RNA Mini Kit manual), a ≥5.9 log10 titer reduction was recorded with no virus recoverable following passages in cell culture.

**Table 3:**
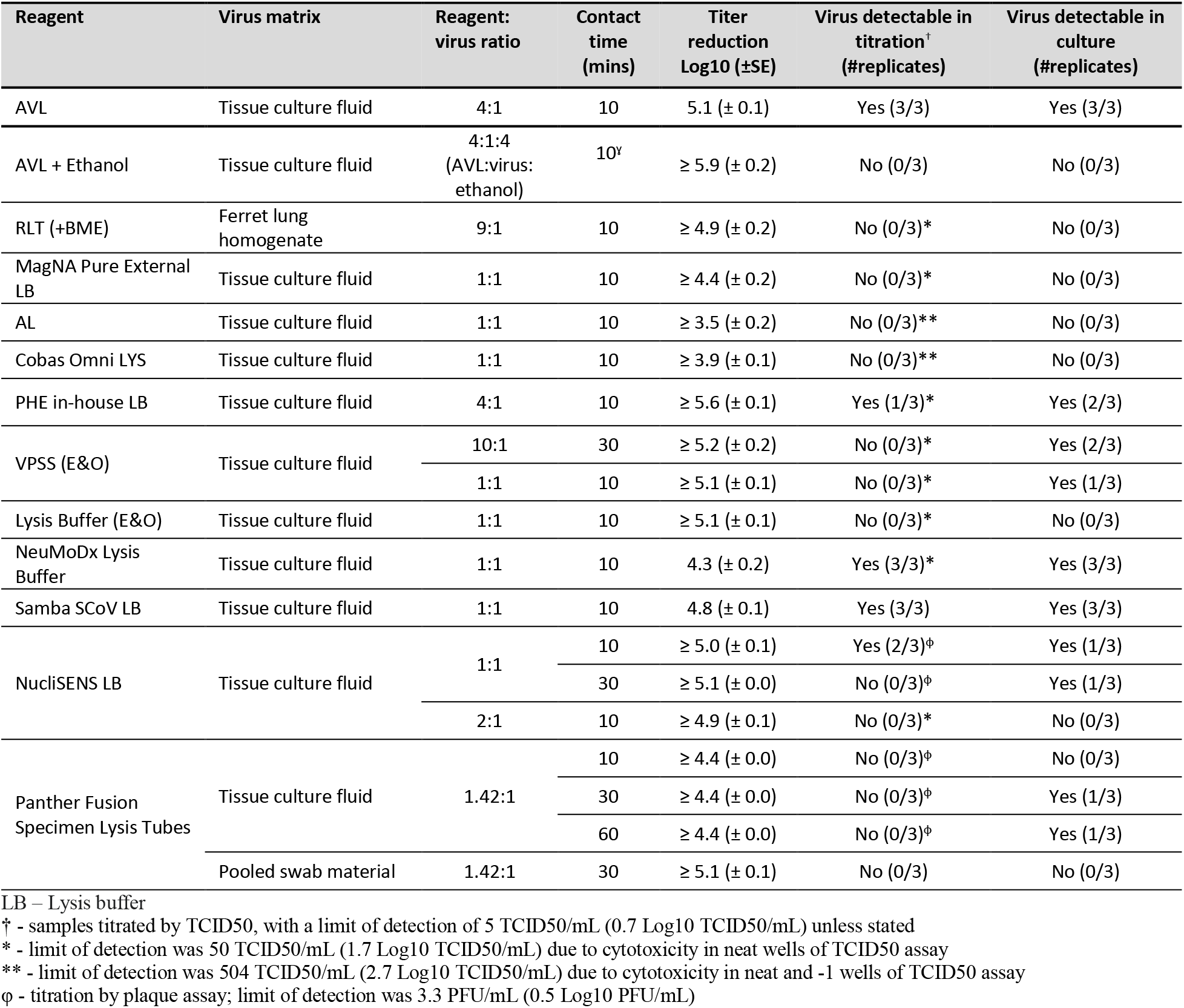
Virus inactivation by molecular extraction reagents

Panther Fusion lysis buffer was further tested against a relevant clinical sample matrix, pooled fluid from oropharyngeal (OP) and nasopharyngeal (NP) swab specimens, resulting in a ≥5.1 log10 titer with no remaining infectious virus detectable. We additionally evaluated the tissue lysis buffer RLT using homogenised ferret lung as sample material, with treatment resulting in a ≥4.8 log10 titer reduction with no residual infectious virus detectable.

### 3.3. SARS-CoV-2 inactivation by detergents

Detergents can be used to inactivate lipid enveloped viruses such as coronaviruses by disrupting the viral envelope, therefore rendering them unable to attach or enter cells (24–27). Here, we evaluated Triton X-100, SDS, NP40 and Tween 20 for their ability to inactivate SARS-CoV-2. SDS treatment at 0.1% or 0.5% reduced titers by ≥5.7 and ≥6.5 log10 TCID50s, respectively, while both concentrations of NP40 reduced titers by ≥6.5 log10 TCID50 with no residual virus detectable following NP40 treatment. In contrast, up to 0.5% Tween 20 had no effect on viral titers. Triton X-100 is commonly used in viral inactivation reagents, and here we show that at both 0.1% and 0.5% v/v concentration, virus titers in tissue culture fluid were reduced by ≥4.9 log10 TCID50s, even with less than 2 min contact time (Table 4). Furthermore, we were unable to recover infectious virus from samples treated with 0.5% Triton X-100 for 10 mins or longer. We also saw effective inactivation of SARS-CoV-2 by SDS, NP40 and Triton X-100 in spiked NP and OP swab specimen fluid, but, importantly, we were not able to replicate this in spiked serum; 1% Triton X-100 only reduced titers in human serum by a maximum of 2 log10 TCID50s with contact times of up to two hours.

**Table 4:**
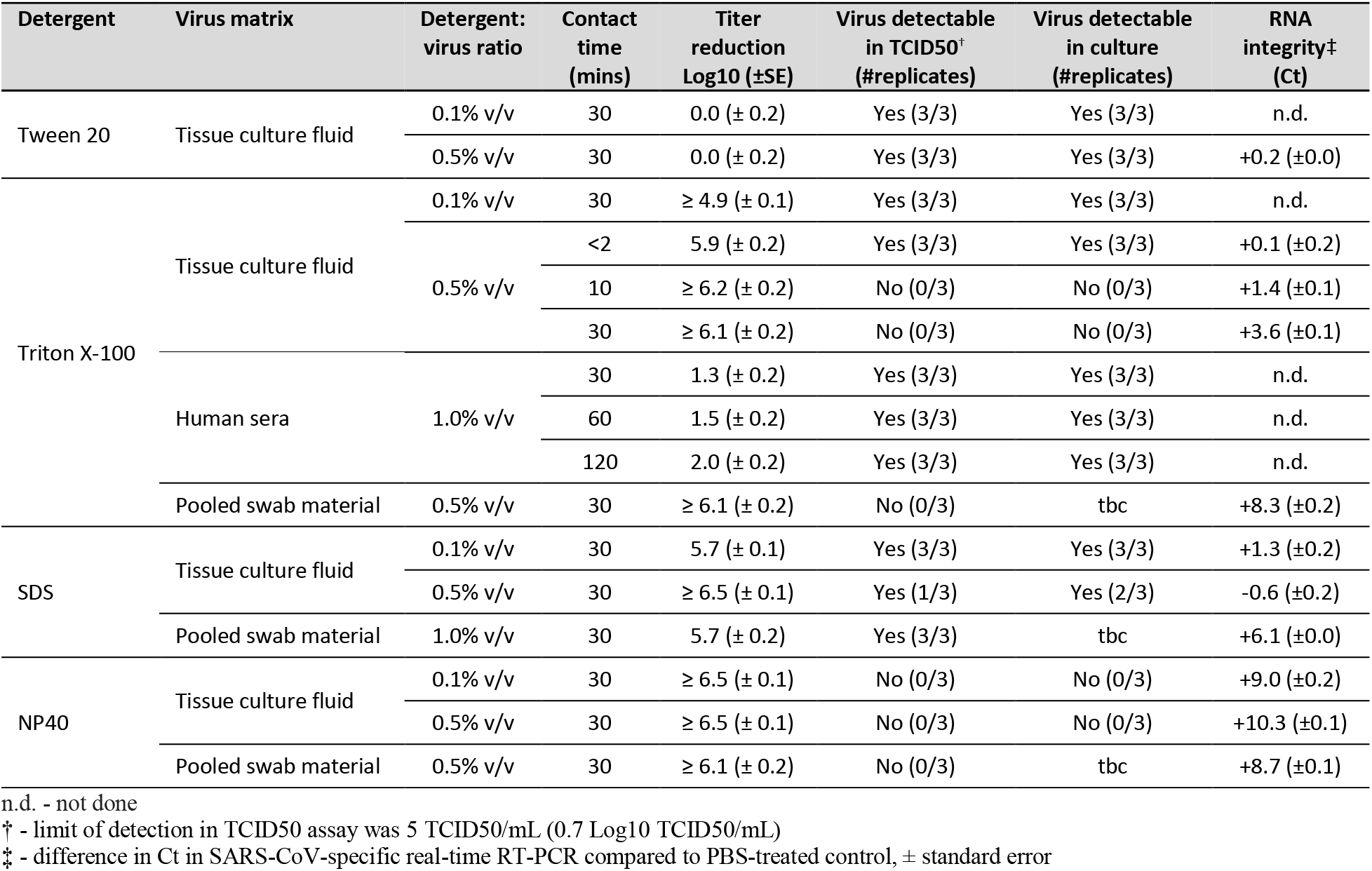
Virus inactivation by detergents

In addition to evaluating inactivation efficacy by detergents, we assessed the effects of treatment on RNA integrity to determine their suitability for inactivation prior to nucleic acid testing. Extracted RNA from treated samples was tested using a SARS-CoV-2-specific qRT-PCR, and the Ct difference between detergent-treated samples and mock-treated controls determined (Table 4). A time-dependent increase in Ct value following treatment with 0.5% Triton X-100 was observed, indicating a detrimental effect on RNA stability with increasing treatment times. Treatment with NP40 had a marked effect, with a 30 minute treatment leading to an increase in 9-10 Cts. While we saw no increase in Ct in tissue culture fluid samples treated with 0.5% SDS, we observed an increase in Ct for SDS-treated swab fluid samples, likely due to an increased concentration of RNases in clinical samples.

### 3.4. SARS-CoV-2 inactivation by other chemical treatments

Fixation and inactivation of viruses by addition of formaldehyde, or a combination of formaldehyde and glutaraldehyde, is a well-established protocol, particularly for diagnostic electron microscopy (28, 29). 4% or 2% formaldehyde treatment for 15 or 60 mins reduced virus titers by ≥4.8 log10 TCID50s when evaluated against a tissue culture fluid matrix, with no remaining infectious virus detectable (Table 5). When infected monolayers were subjected to the same treatment protocol, titer reductions were all ≥6.8 log10 TCID50s, with 60 min contact time moderately more effective than 15 min. However, in this format, a 60 min 4% formaldehyde treatment was the only one from which no infectious virus was detectable. A mixture of 2% formaldehyde with 1.5% glutaraldehyde tested on infected monolayers reduced virus titers by ≥6.7 log10 TCID50s with no remaining infectious virus detectable for both a 15 and 60 min contact time. Polyhexanide biguanide (PHMB) is a polymer used as a disinfectant and antiseptic, evaluated here as a potential lysis buffer, but it was only able to reduce viral titers by 1.6 log10 TCID50s at the highest concentration tested (2%).

**Table 5:**
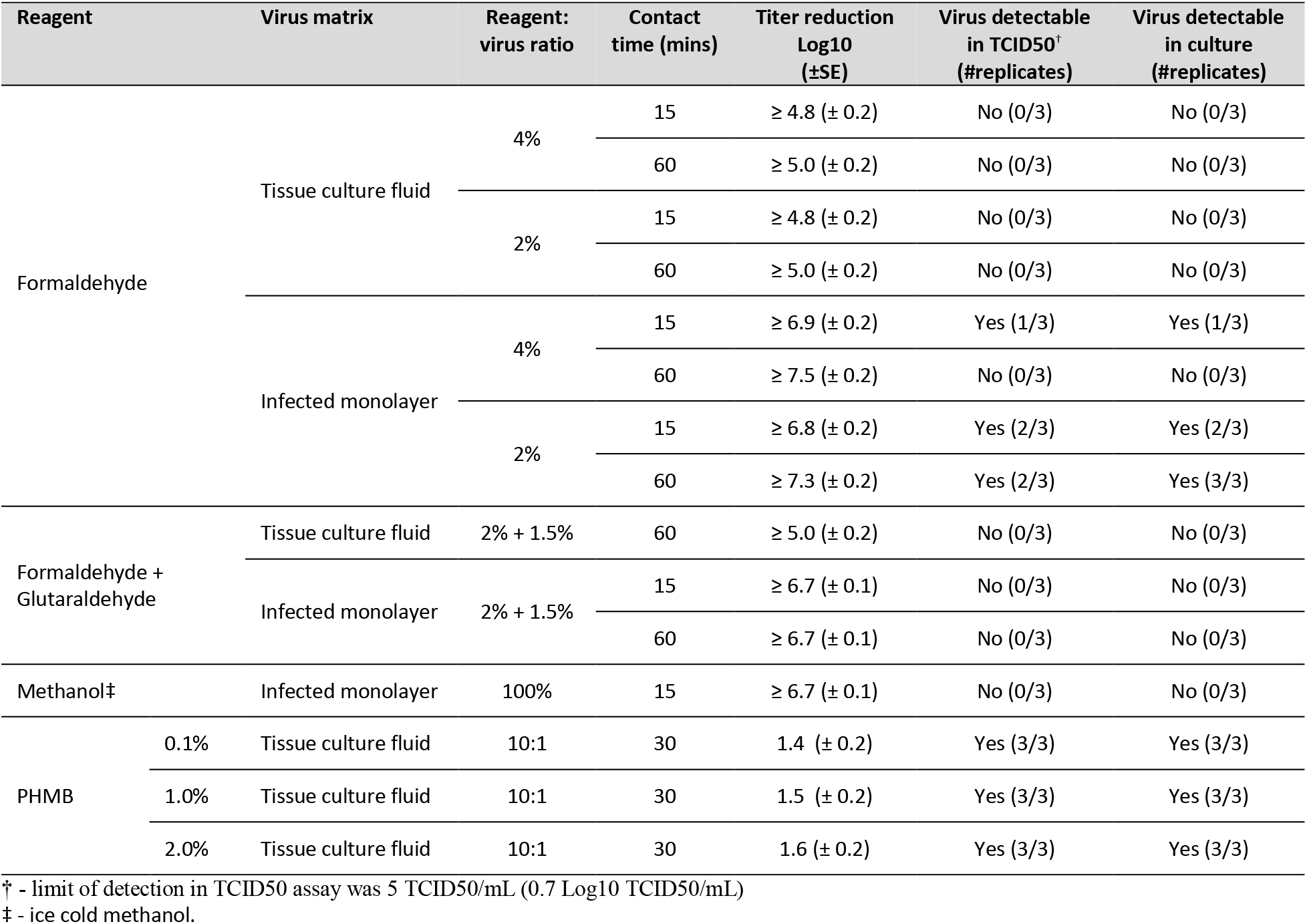
Other reagent types

## 4. Discussion

Samples containing infectious SARS-CoV-2 require an initial inactivation step before downstream processing; given the rapid emergence of SARS-CoV-2, these inactivation protocols have been guided by existing data for other coronaviruses and there is an urgent need to both confirm these historical data using the new virus and to validate new approaches for inactivating SARS-CoV-2. We therefore analysed numerous commercially and commonly available reagents used by public health agencies and research laboratories around the world in their response to the pandemic. In addition, to address challenges of reagent cytotoxicity in inactivation evaluation, we provide data on the effectiveness of filtration methods for removing cytotoxicity from chemically treated samples.

Knowledge of the expected amount of infectious virus in clinical samples obtained from COVID-19 patients is important when applying viral inactivation study data to diagnostic sample processing, allowing end users to interpret whether material they are handling is likely to represent an infectious risk to themselves and others. These values are dependent on several factors, including time post symptom onset, duration of symptoms, time elapsed between sampling and testing, the presence of neutralizing antibody responses, and immunocompetency of the individual (30). Data regarding quantitative infectious viral levels in typical clinical specimens are minimal, with most studies reporting viral loads determined by qRT-PCR only (31–33). One study of 90 qRT-PCR positive NP or endotracheal (ETT) samples from COVID-19 patients estimated the median titer at 3.3 log10 TCID50/mL (30). Given we demonstrate >4 log_10_ reduction in titer for all specimen transport reagents, this suggests that these reagents may considerably decrease, even eliminate, the infectivity of a clinical sample. However, our observation that residual virus could be recovered from most treated samples indicates that these media cannot be assumed to completely inactivate SARS-CoV-2 in samples and that additional precautionary measures should be taken in laboratories when it comes to sample handling and transport.

Limited SARS-CoV-2 inactivation data on molecular extraction reagents used in nucleic acid detection assays are currently available. One study reported that Buffer AVL either alone or in combination with ethanol was not effective at completely inactivating SARS-CoV-2 (15). By contrast, we could not recover any infectious virus from samples treated with AVL plus ethanol, consistent with previous studies indicating that AVL and ethanol in combination is effective at inactivating MERS and other enveloped viruses (10, 34), and indicating that both AVL and ethanol steps of manual extraction procedures should be performed before removal of samples from primary containment for additional assurance. Our detergent inactivation data, indicating that SDS, Triton X-100 and NP40, but not Tween 20, can effectively inactivate SARS-CoV-2 in tissue culture fluid and in pooled NP and OP swab fluid, corroborate findings of a recent study (17); however, as has been demonstrated for other viruses (31), we observed an inhibitory effect of serum on virus inactivation by detergent, highlighting the importance of validating inactivation methods with different sample types.

Based on our findings comparing filtration matrices, we found that the optimum method for reagent removal for inactivation studies is determined by evaluating three factors: (i) effectiveness of cytotoxicity removal; (ii) efficiency of virus recovery; and (iii) the ease of performing these methods within a containment space. Methods permitting complete removal of cytotoxic reagent components with no or little effect on virus recovery give assurance that low levels of residual virus, if present, could be detected in virus inactivation studies. During reagent testing, there were several instances where we noted residual cytotoxicity in the neat eluate contrary to what was expected based on the initial reagent removal data and is likely due to the extended incubation period required for inactivation testing (up to 7 days, compared with overnight for cytotoxicity evaluation). In all cases however, we were still able to enhance the levels of titer reduction detectable when compared with what would have been achieved by sample dilution alone.

In conclusion, we have evaluated methods for straightforward, rapid determination of purification options for any reagent prior to inactivation testing, enabling establishment of effective methods for sample purification while minimising virus loss. This is applicable to inactivation studies for all viruses (known and novel), not only SARS-CoV-2. We have applied these methods to obtain SARS-CoV-2 inactivation data for a wide range of reagents in use (or proposed for use) in SARS-CoV-2 diagnostic and research laboratories. In addition to guiding laboratory risk assessments, this information enables laboratories to assess alternative reagents that may be used for virus inactivation and nucleic acid extraction, particularly considering concerns about extraction reagent availability due to increased global demand caused by the COVID-19 pandemic. Furthermore, chemical treatments evaluated here are commonly used for inactivation of a wide range of different viruses and other pathogens, and the results presented may be used to directly inform and improve the design of future inactivation studies.

## Acknowledgments

The authors would like to thank: The Respiratory Virus Unit at PHE Colindale, and the Virology Laboratories at PHE Cambridge and PHE Bristol for donation of pooled respiratory samples; and Ayoub Saei at the Statistics Unit, PHE Colindale for statistical advice.

The views expressed in this article are those of the author(s) and are not necessarily those of Public Health England or the Department of Health and Social Care

---

**Supplementary Table 1:** Reagent Details

| Reagent Type | Reagent | Manufacturer Cat# | Reagent composition | Recommended ratio of sample to reagent | Recommended contact time |
| --- | --- | --- | --- | --- | --- |
| Specimen Transport Tube Reagents | Virus Transport and Preservation Medium (Inactivated) | BioComma Ltd. #YMJ-E | Not known | Swab placed directly into tube containing 3mL reagent | None given |
|  | Sigma MM | Medical Wire #MWMM | Guanidine thiocyanate, Ethanol (concentrations unknown) | Up to 1 vol sample to 1.5 vols reagent (up to 0.67:1) | None given |
|  | eNAT | Copan #608CS01R | 42.5-45% guanidine thiocyanate, detergent, Tris-EDTA, HEPES. | Swab placed directly into tube containing 1 or 2mL reagent. For urine, 3:1 | None given |
|  | Primestore | Longhorn #PS-MTM-3 | <50% guanidine thiocyanate, <23% ethanol | 1:3 | None given |
|  | Cobas PCR | Roche #08042969001 | ≤40% guanidine hydrochloride, Tris-HCl | Swab placed directly into tube | None given |
|  | Aptima Specimen Transport Medium | Hologic #PRD-03546 | Not known | Swab OR 0.5mL VTM sample added to tube containing 2.9mL buffer | None given |
|  | DNA/RNA Shield | Zymo Research #R1100 | Not known | 1:3 | None given |
|  | 40% GHCL | Oxoid/Thermo Fisher #EB1351A | 28.3% guanidine hydrochloride, 2.1% Triton X-100, Tris-EDTA | Swab placed directly into tube | None given |
|  | 2M GITC | Oxoid/Thermo Fisher #EB1349A | 18.9% guanidine thiocyanate, 2.4% Triton X-100, Tris-EDTA | Swab placed directly into tube | None given |
|  | 4M GITC | Oxoid/Thermo Fisher #EB1350A | 31.8% guanidine thiocyanate, 2.0% Triton X-100, Tris-EDTA | Swab placed directly into tube | None given |
| Molecular Extraction Reagents | NucliSENS Lysis Buffer | Biomerieux #200292 | 50% guanidine thiocyanate, <2% Triton X-100, <1% EDTA | 1:2-1:200 | 10 mins |
|  | Panther Fusion | Hologic #PRD-04339 | Not known | 1:1.42 |  |
|  | Buffer AVL | QIAGEN #19073 | 50-70% guanidine thiocyanate | 1:4 | 10 mins |
|  | MagNA Pure 96 External Lysis Buffer | Roche #06374913001 | 30-50% guanidine thiocyanate, 20-25% Triton X-100, <100mM Tris-HCl, 0.01% bromophenol blue. | 1:1 | None given |
|  | Buffer AL | QIAGEN #19075 | 30-50% guanidine hydrochloride, 0.1-1% maleic acid | 1:1 | None given |
|  | Cobas Omni LYS | Roche #06997538190 | 30-50% guanidine thiocyanate, 3-5% dodecyl alcohol, ethoxylated, 1-2.5% dithiothreitol | No instructions for use as off-board lysis buffer | None available |
|  | L6 (Kingfisher formulation) | PHE Media Services | 96.6% guanidine thiocyanate, 1.9% Triton X-100, Tris-EDTA | None available | None available |
|  | Buffer RLT | QIAGEN #79216 | 30-50% guanidine thiocyanate | Tissue to be homogenized directly in undiluted buffer | None given |
|  | NeuMoDx Viral Lysis Buffer | NeuMoDx Molecular, Inc. #401600 | <50% guanidine hydrochloride, <5% Tween 20, <1% EDTA, <0.1% sodium azide | 1:1 | None given |
|  | VPSS | E&O Laboratories #BM1675 | Not known | Not known | Not known |
|  | Lysis Buffer | E&O Laboratories #BM1676 | Not known | Not known | Not known |

**Supplementary Figure 1:**
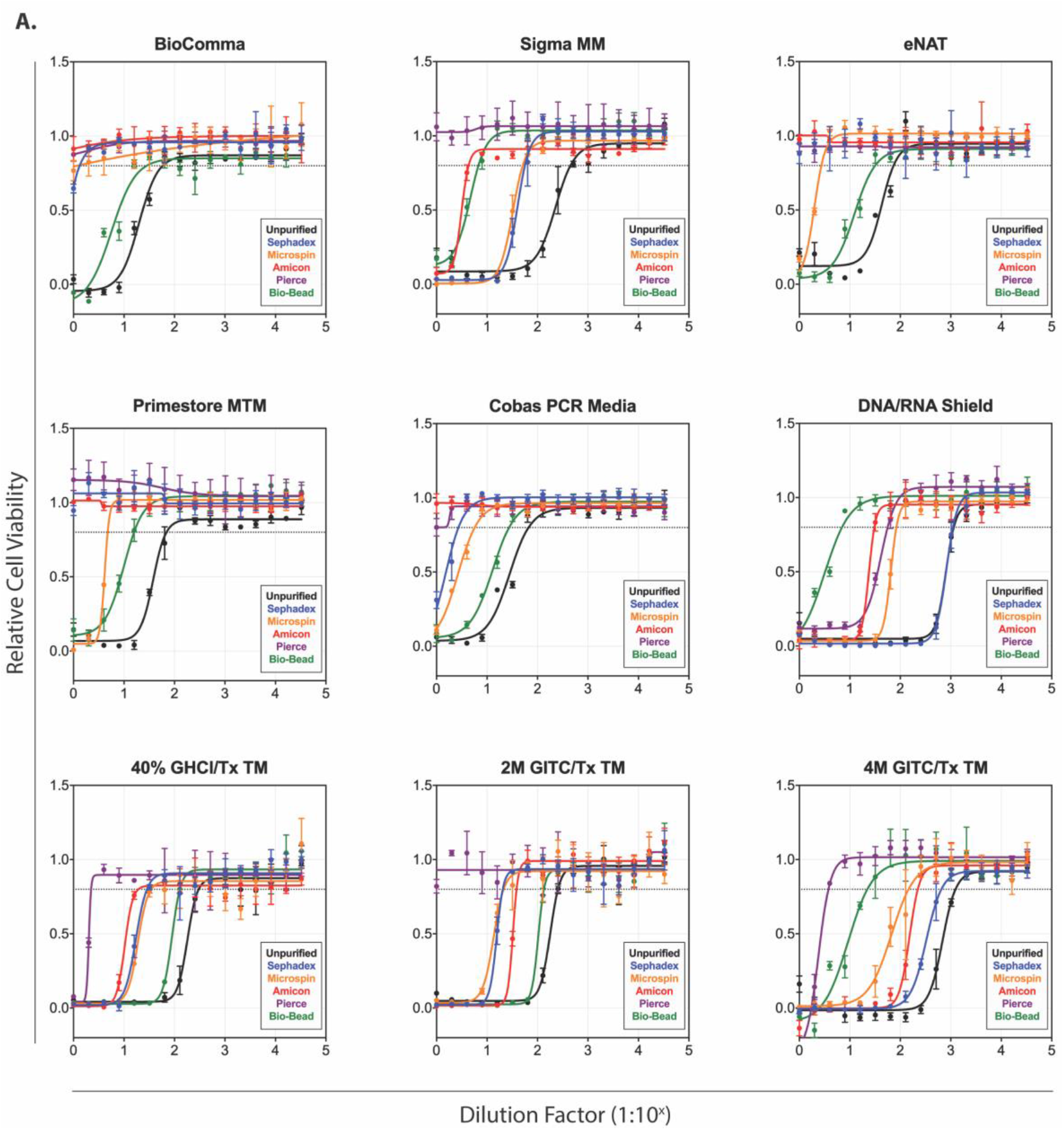
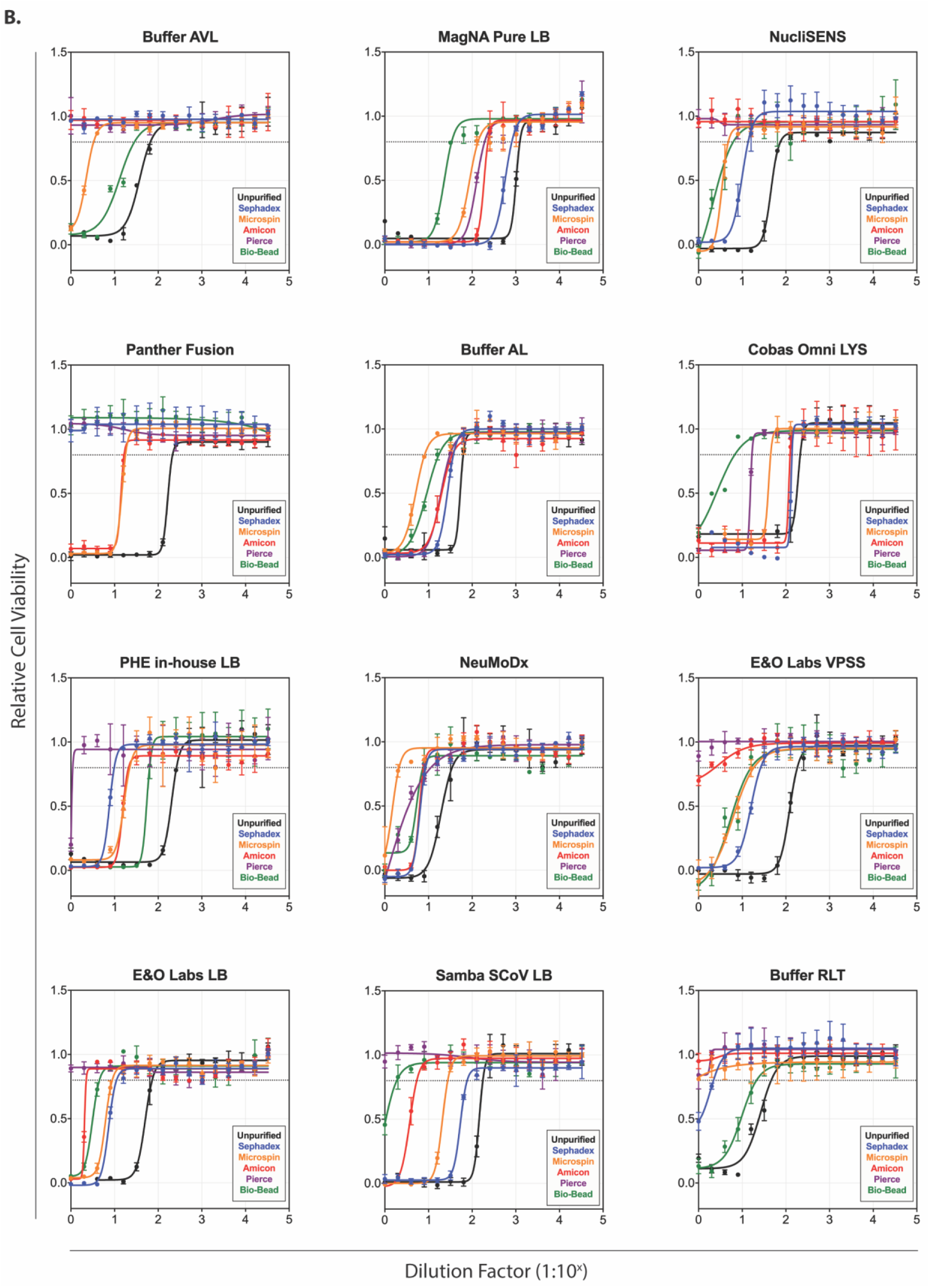
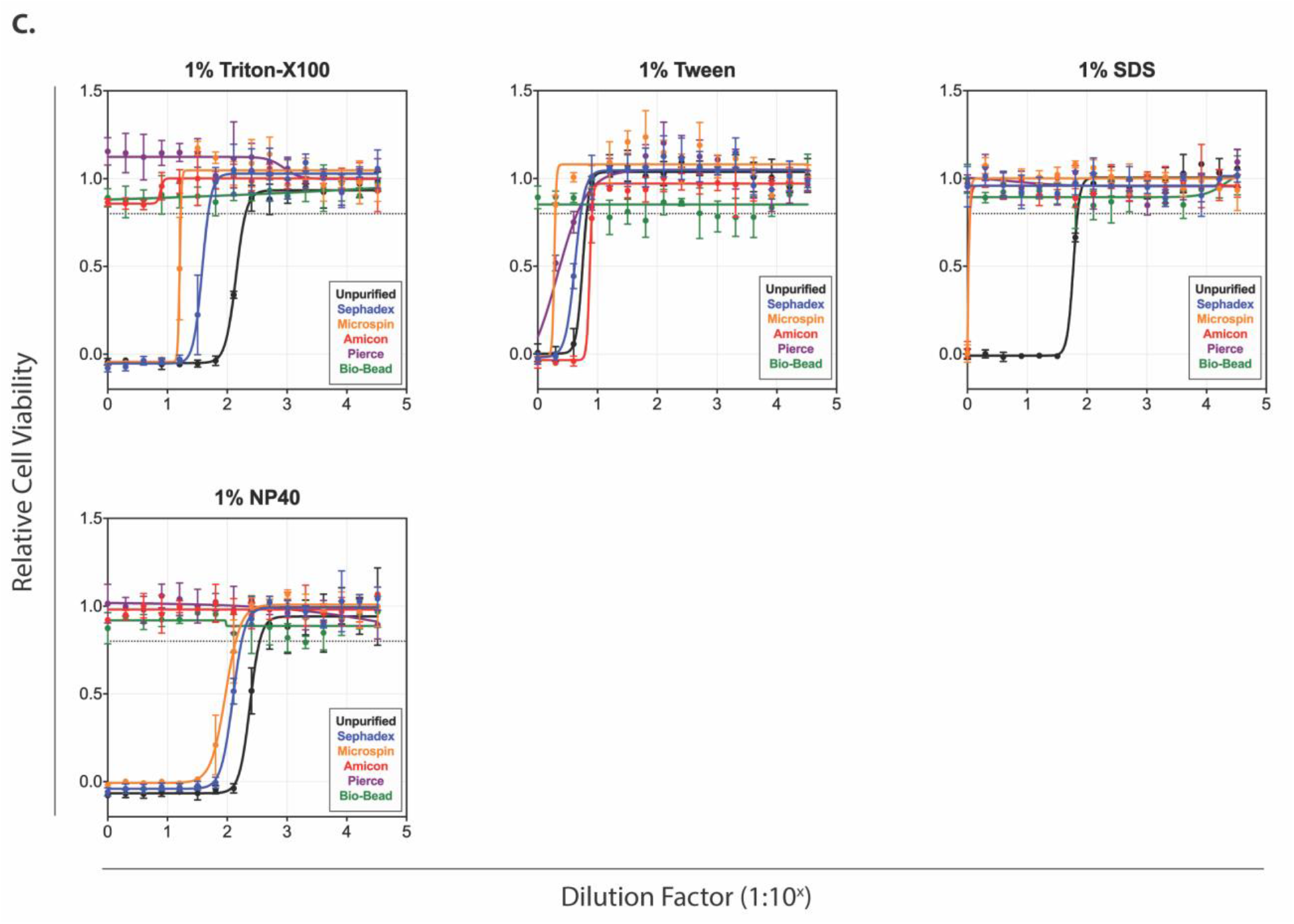
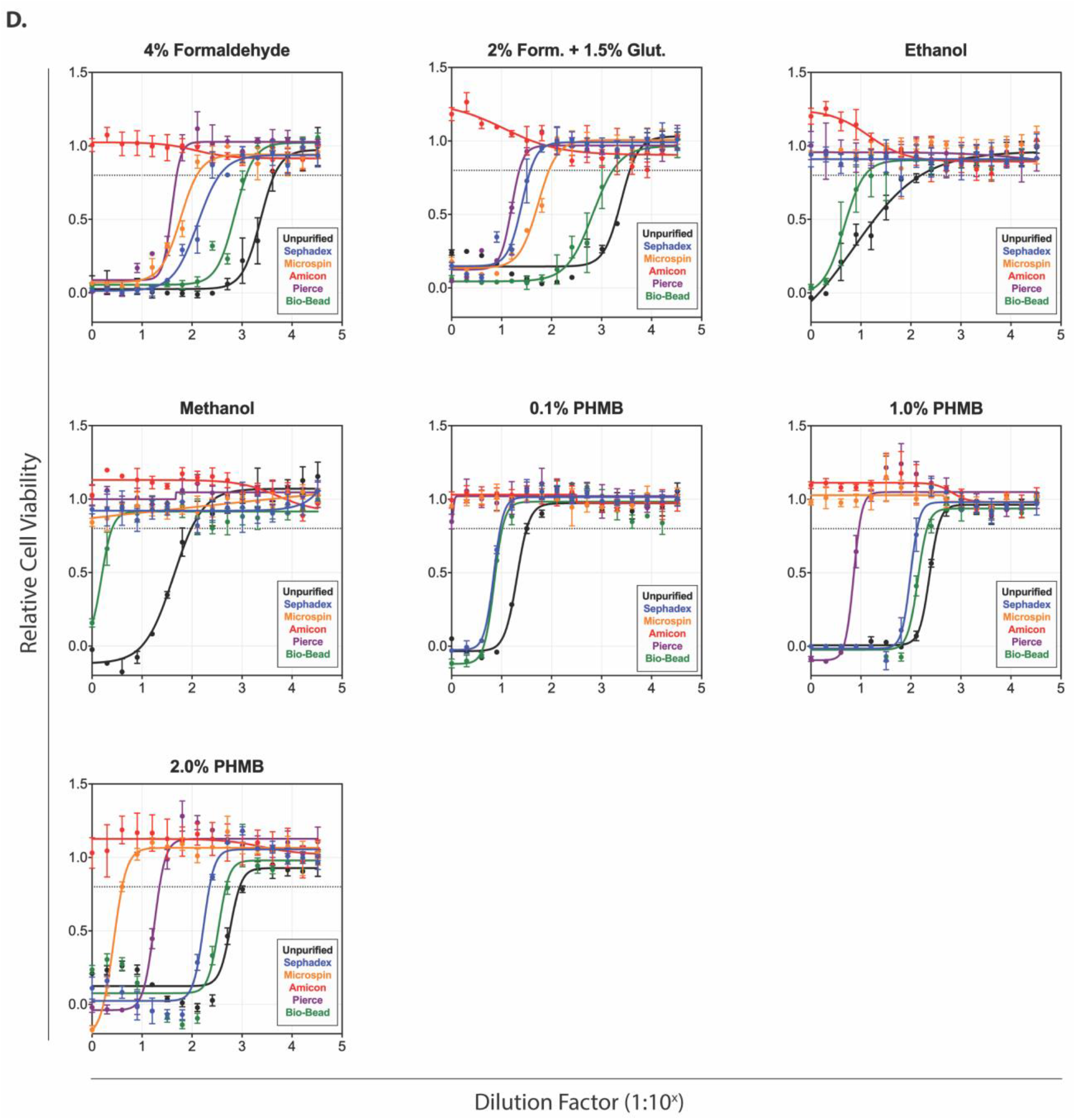
Cytotoxicity of virus inactivation reagents after passing through purification matrices. Concentration-response curves in Vero cells treated with a 2-fold serial dilution of reagent. At 24 h post treatment cell viability was determined, with values normalized to mock treated cells. Each point represents the mean of triplicate wells, with error bars indicating standard deviation. Graphs are representative of at least 2 independent experiments. Matrices used: Sephadex LH-20 (blue); Sephacryl S400HR (orange); Amicon Ultra 50kDa molecular weight cut off (red); Pierce detergent removal spin columns (DRSC) (purple); or Bio-Bead SM2 (green). **(A)** Reagents used in specimen transport tubes: GHCl - guanidine hydrochloride; GITC - guanidinium isothiocyanate; Tx – Triton X-100; TM – Transport Medium **(B)** Reagents used in molecular extraction protocols: PHMB - polyhexamethylene biguanide. **(C)** Detergents commonly used for virus inactivation: SDS - sodium dodecyl sulfate; NP40 - nonyl phenoxypolyethoxylethanol. **(D)** Other reagents commonly used for virus inactivation.

